# Diet gel-based oral drug delivery system for controlled dosing of small molecules for microglia depletion and inducible Cre recombination in mice

**DOI:** 10.1101/2025.01.23.634530

**Authors:** Joel Jovanovic, Megan L. Stone, Samantha R. Dooyema, Yuankai K. Tao, Sabine Fuhrmann, Edward M. Levine

## Abstract

Small molecules like PLX5622 for microglia depletion and Tamoxifen for inducible Cre recombination are commonly used in mouse research. Traditional application methods, such as chow or oral gavage and injections, have limitations, including uncontrolled dosage and risk of injury. To address this issue, we have developed an alternative oral drug delivery system using a gel-based rodent maintenance diet that allows for controlled consumption and adjustment of dosage and is suitable for water-insoluble small molecules. We tested DietGel^®^ 93M (93M) infused with PLX5622 (0.8 mg/g and 2.0 mg/g) in the *Cx3cr1*^*gfp/+*^ retinal microglia reporter mouse and Tamoxifen-infused 93M (0.3125 mg/g) in the Rlbp1-Cre^ERT2^;*Rosa*^*ai14*^ mouse with an inducible tdTomato reporter in retinal Müller glia. Mice were single-caged and received daily batches of PLX5622-infused 93M over 14 days or Tamoxifen-infused 93M for one or three days followed by a 14-day observation period. Longitudinal scanning laser ophthalmoscopy *in vivo* and fixed tissue imaging were used to track GFP and tdTomato expression. Following evaluation of a suitable 93M consumption rate (g/d) to sustain body weight, the PLX5622-93M diet at both concentrations showed a 94% microglia depletion rate at 3 days and >99% after one and two weeks. The Tamoxifen-93M diet confirmed suitability for inducible Cre recombination, with significant treatment-time dependent efficacy and a positive correlation between total Tamoxifen dose and tdTomato expression. This study demonstrates that a diet gel-based drug delivery system offers a controllable and less invasive alternative to current drug application methods for PLX5622 and Tamoxifen.

## Introduction

Over the past years, different drug delivery systems for rodents have been established in biomedical research. The preferred route of delivery often depends on the drug’s properties, the drug vehicle, the required dose, and the rodent’s condition ^1,2^. Among the most used systemic delivery methods using the oral route are gavage, drinking water or hydration gels, and drug-infused chow, while other drugs require bypassing the first pass effect by using the parenteral route, such as intravenous (tail vein), subcutaneous or intraperitoneal (i.p.) injections ^1,3–6^. Even though these systems are widely used, they have disadvantages that must be considered such as the risk of injury, stress to the animal, and control over dosage.

Recently, a broad range of small molecules became available for a variety of purposes ^7,8^. Among commonly used small molecules in research are PLX5622 for microglia depletion and Tamoxifen for CreER recombination ^9–14^. PLX5622 is a small molecule kinase inhibitor that selectively blocks the CSF1R signaling required for cell survival in monocytes, and in microglia and macrophages in the CNS ^15^. Tamoxifen is a selective estrogen receptor modulator widely used in combination with a Cre recombinase-LoxP system for tissue-specific and temporal gene regulation in mice ^16–18^.

Due to the water-insoluble properties of both drugs, the standard oral application methods for these small molecules include drug-infused chow (ad libitum) for PLX5622 and Tamoxifen or oral gavage for Tamoxifen. However, these methods may not always be suitable. While chow lacks control over dosage (mg/g) and consumption rate (g/day) between rodents, oral gavage risks injury and stress to the animals. Therefore, an alternative that mitigates these effects is needed. In this study, we have developed a diet gel-based oral drug delivery system that allows for dosage adjustment and is suitable for water-insoluble small molecules using the complete gel-based maintenance diet ClearH_2_O^®^ DietGel^®^ 93M (93M).

## Results

### A 93M DietGel^®^ consumption rate of 8 g/day allows for complete consumption and steady body weight

To facilitate the transition from regular chow to diet gel and to allow for a time-matched start of the experiment across all conditions, mice were fasted for 16 hours before the experimental start. Body weight was tracked throughout the experiments with the post-fast body weight serving as the baseline. This allowed us to directly assess the effect of the diet gel with and without drug infusion.

As a first step, we determined the optimal daily consumption rate of drug-free 93M by assigning three feeding groups to receive either 6, 8, or 10 grams per day (g/d) of 93M over 14 days (**Fig. 1a**). This experiment used B6129SF1/J mice which share the genetic background with the two strains used for the following drug-incorporation studies. Body weight and the residual amount of food (unconsumed 93M) were measured daily after each feeding cycle, including a body weight measurement right before and after an initial fasting period of 16 hours before the 93M feeding start. Each feeding group included 3 male and 3 female mice, and all mice were single caged for the experiments.

**Fig. 1:**
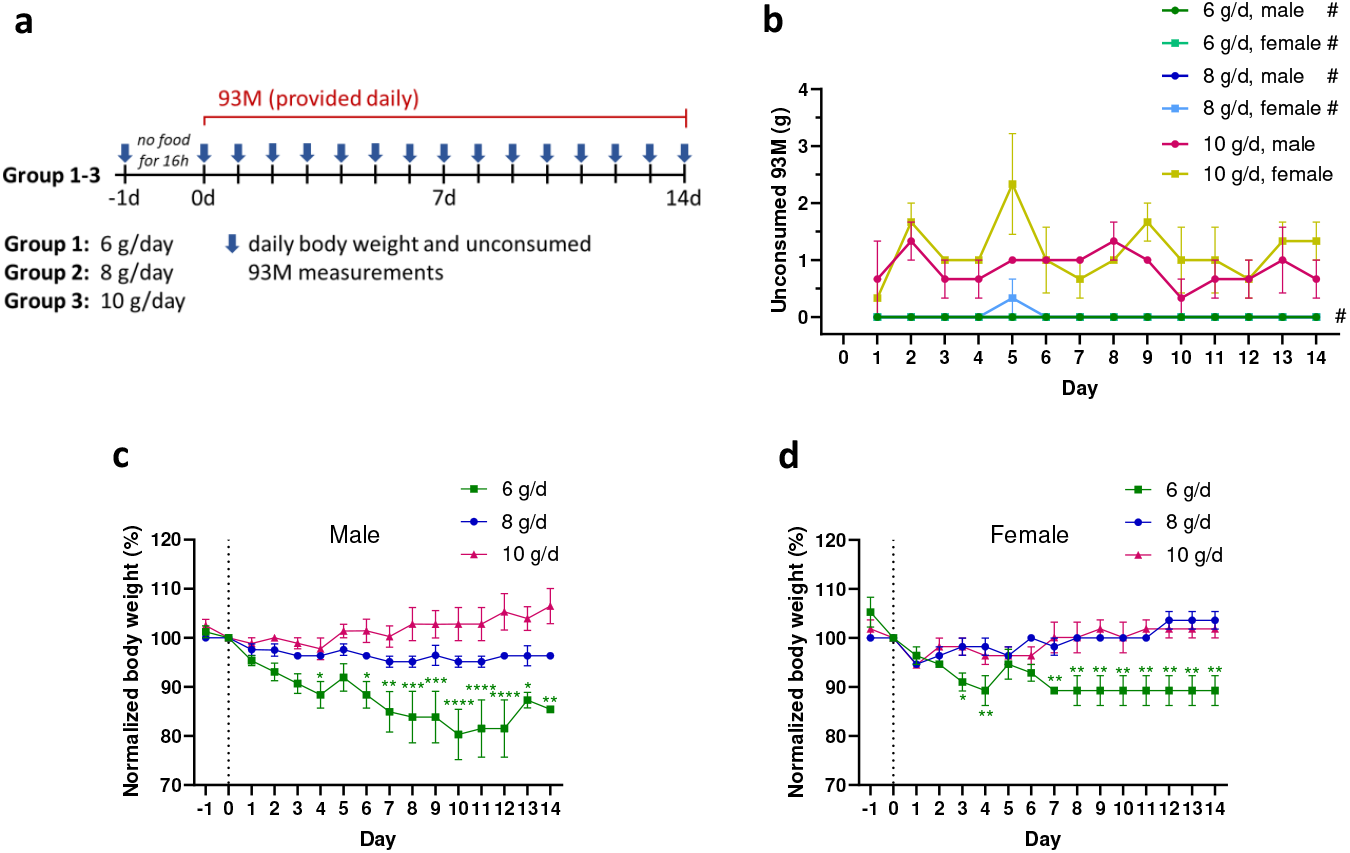
Consumption rate determination for steady body weight and complete consumption of provided 93M in B6129SF1/J mice. **a**, Experimental design of feeding groups receiving either 6g, 8g or 10g of drug-free 93M every 24 hours over 14 days. Unconsumed 93M (g) and body weight (g) were measured after each 24h-feeding cycle, including a body weight measurement before (day -1) and after (day 0) an initial fasting period of 16 hours. **b**, Unconsumed 93M (g) after each feeding cycle per feeding group and sex. #: The data points are at 0 g of unconsumed 93M. **c**,**d**, Temporal body weight measurements normalized and compared to the post-fasting body weight (day 0) in males and females. N = 6 mice (3 males, 3 females) per feeding group. Statistics: Two-way ANOVA with Dunnett’s multiple comparisons tests for (**c**,**d**). Results are shown as Mean ± SEM. Significance levels: \**P* < .05, \*\**P* < .01, \*\*\**P* < .001, \*\*\*\**P* < .0001.

**Fig. 1b** shows the mean amount of unconsumed food at each time point for males and females. For both sexes, 6 and 8 g were completely consumed over the course of the experiment except for one female on day 5. In contrast, unconsumed food was left behind every day with 10 g/d. This shows that males and females consistently consume 6 or 8 g/d and leave food behind when given 10g/d with an average of 0.86 ± 0.09 g/d for males and 1.14 ± 0.12 g/d for females without a significant difference between them (*P* = 0.0588, two-tailed unpaired t-test).

A potential confound that could underestimate the amount of unconsumed food and lead to miscalculation of an ingested drug dose is evaporation. Our temporal weight measuring data of 93M at 6 g, 8 g, and 10 g held in ventilated cages show mass reductions between 25% to 40% over 24 hours, depending on the initial amount of 93M and on the drug-vehicle mix-in (**Supplementary Fig. 1**). However, we observed that mice in the 6 g/d and 8 g/d groups did not leave food residues in the petri dishes, indicating that they consumed the full portions. For the 10 g/d group, however, the true mass of the unconsumed 93M could be greater than what was measured due to the uncertainty of when the food was consumed over a 24-hour period.

To determine the effect of the three different provided amounts of 93M on body weight maintenance, animals were weighed daily, and the measurements were normalized to the initial post-fasting weights for each animal (day 0) (**Fig. 1c,d**). Two-way ANOVA shows that both sexes have significant variations in body weight changes depending on the feeding amount (*p*_males_<0.0001, *p*_females_<0.0001) and time (*p*_males_=0.0036, *p*_females_=0.0013). These significant changes were specifically observed in mice given 6 g/d which caused significant drops in body weights, with one male mouse removed from the study on day 12 due to a weight loss greater than 30%.

Overall, both males and females fed 8 g/d of 93M showed a steady body weight over time and completely consumed the daily portions. Therefore, we decided to use 8 g/d of 93M for the following drug incorporation studies.

### Efficient retinal microglia depletion kinetics in mice treated with low and high dose PLX5622-infused 93M DietGel^®^

The feasibility and efficacy of PLX5622-infused 93M (PLX5622-93M) treatment were tested in *Cx3cr1*^*gfp/+*^ microglia reporter mice split into a low (0.8 mg/g) and a high (2.0 mg/g) dosage group with feedings of 8 g/d over 14 days (**Fig. 2a**). Body weights and unconsumed food were measured daily. Both groups included 8 mice with 4 males and 4 females. Scanning laser ophthalmoscopy (SLO) was used to visualize the microglia depletion kinetics *in vivo* at day 0 (pre-treatment baseline), 3, 7, and day 14. Endpoint histology in retinal whole mounts was used to confirm the *in vivo* results.

**Fig. 2:**
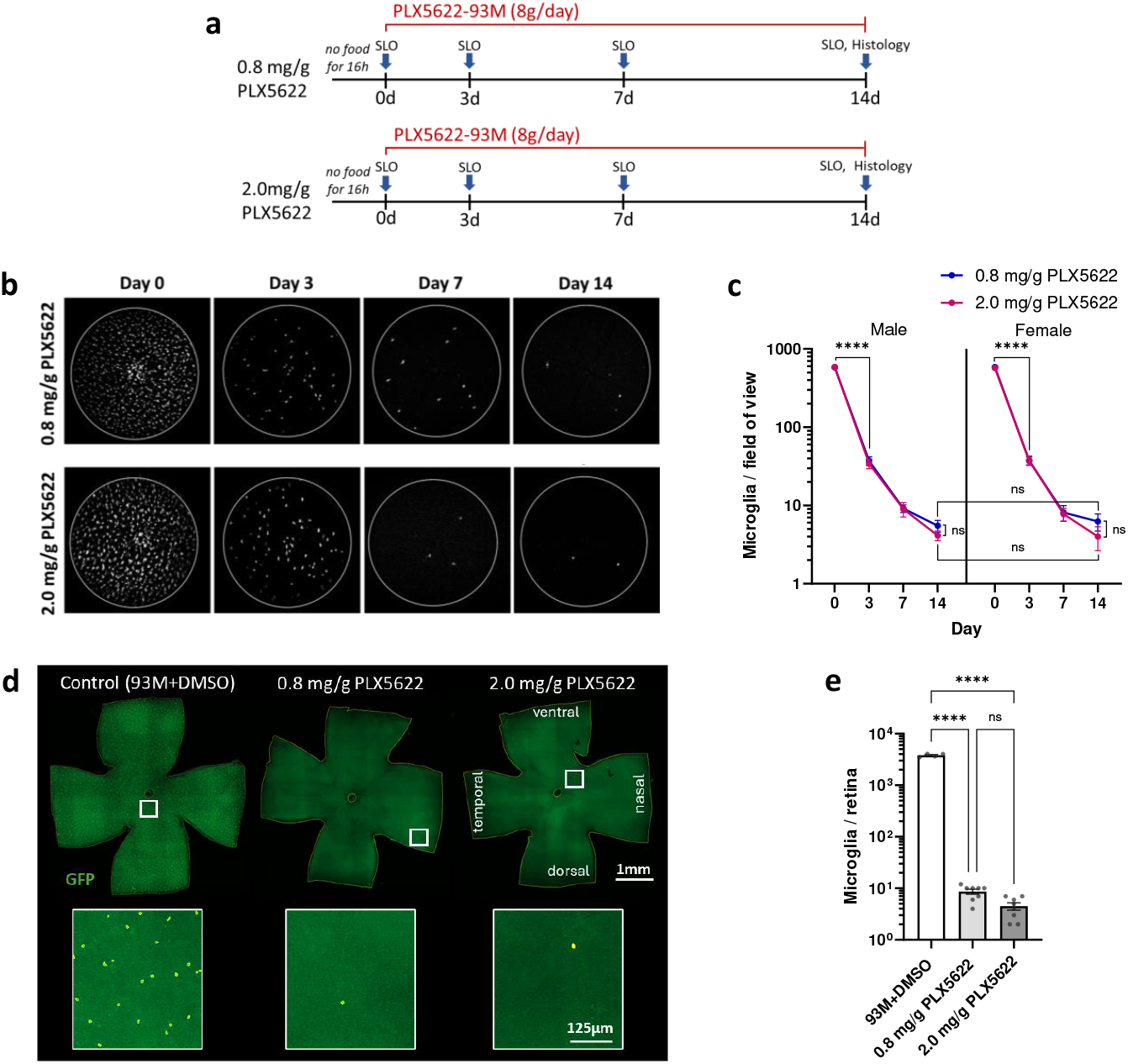
Feasibility and efficacy evaluation of PLX5622-infused 93M for retinal microglia depletion in GFP expressing microglia reporter mice. **a**, Experimental design with two PLX5622-93M dosage groups (0.8 mg/g and 2.0 mg/g) in heterozygous B6.129P2(Cg)-*Cx3cr1* ^*tm1Litt*^/J mice receiving 8 g/d over 14 days, including daily body weight and unconsumed food measurements. SLO was performed on day 0, 3, 7, and 14. **b**, Representative longitudinal SLO image series per treatment group shows retinal microglia depletion *in vivo*. **c**, Quantification of microglia counts in SLO images per sex and dosage group. **d**, Representative endpoint histology (day 14) of whole mounted retina per dosage group, including a control retina treated with DMSO-infused 93M. The inset shows a magnified section with masked microglia (yellow boundary) for better visualization. **e**, Quantification of microglia counts in retinal whole mounts over both sexes per dosage group. N = 8 mice (4 males, 4 females) per PLX5622 dosage group. Statistics: Two-way ANOVA with Tukey’s multiple comparisons test for (**c**) and one-way ANOVA with Tukey’s multiple comparisons test for (**e**). The results are shown as Mean ± SEM. Significance level: \*\*\*\**P* < .0001.

SLO proved the feasibility of 93M as a drug delivery system for PLX5622, as evidenced by the loss of GFP+ cells, the readout for microglia depletion (**Fig. 2b**). Surprisingly, both dosages showed significant depletion by 3 days of treatment (*P* < 0.0001) with similar depletion kinetics for both sexes (**Fig. 2c**). Animals treated with 0.8 mg/g showed a 93.6% reduction in microglia at day 3, 98.5% at day 7, and 99.1% at day 14. Animals treated with 2.0 mg/g showed a 93.8% reduction in microglia at day 3, 98.6% at day 7, and 99.3% at day 14.

Quantification of fixed retinal whole mount tissues on day 14 confirms these findings (**Fig. 2d,e**), with a depletion of 99.8% for the low dose and 99.9% for the high dose when compared to the vehicle-treated control group. In alignment with the SLO findings on day 14, there is no significant difference between both dosages over both sexes with an adjusted *P*-value of 0.9973, while both have a significant effect in comparison to the DMSO-93M control group (*P* < 0.0001) on day 14.

As expected, the consumption of the PLX5633-93M was complete throughout the experiment, except for the first two feeding cycles in females (**Supplementary Fig. 2a**). Interestingly, the females that received either dosage and the males treated with the low dose regained their initial body weight over the treatment course, while the males treated with 2.0 mg/g stayed at the level of the post-fasting weight (**Supplementary Fig. 2b,c**). Consistent with this, the males treated with the higher dose had significantly lower normalized body weights from day 4 onwards compared to the other treatment groups (**Supplementary Fig. 2d**). These observations reveal a specific dose-dependent effect of PLX5622 in male mice that extends beyond microglia depletion (see Discussion).

To exclude potential effects on the microglia counts caused by the drug vehicle DMSO in the 93M, a separate SLO imaging session was conducted with 93M infused with DMSO only (**Supplementary Fig. 2e**). The mice received 8 g daily feedings of DMSO-infused 93M and were imaged at day 0 (pre-treatment) and on days 7 and 14 of the treatment. The microglia counts in the *in vivo* images show in comparison to the pre-treatment condition (578.50 ± 11.49) no effect on the total number of microglia present on day 7 (573.25 ± 19.77; *P* = 0.9963, one-way ANOVA with Dunnett’s multiple comparisons test) and day 14 (586.75 ± 13.92; *P* = 0.9154, one-way ANOVA with Dunnett’s multiple comparisons test).

### Tamoxifen-infused 93M DietGel^®^ treated mice successfully induce tdTomato expression in retinal Müller glia

To test for feasibility and control over the level of Cre recombination using Tamoxifen-infused 93M (Tamoxifen-93M), we used Rlbp1-Cre^ERT2^;*Rosa*^*ai14*^ mice, which express tdTomato in Müller glia after Tamoxifen treatment ^19–21^. Two treatment groups were designed to test for different Tamoxifen exposure times: one with a short (1-day treatment) and one with a long exposure (3-day treatment) (**Fig. 3a**). All mice received 8 g of Tamoxifen-infused 93M per day of treatment with a Tamoxifen concentration in the food of 312.5 μg/g. This daily dose is approximately equivalent to a single dose of 100 μg/g body weight by oral gavage in a mouse that weighs 25 g. Each group included 8 mice with 4 males and 4 females. Longitudinal SLO imaging was performed to detect Cre-dependent tdTomato expression induction in retinal Müller glia. Baseline expression levels were established before treatment began. Both groups were followed up 7- and 14-days post-Tamoxifen treatment, including a 1-day post-treatment time point for the 1-day treatment group. Endpoint histology was used to confirm the tdTomato expression levels in fixed retinal whole mounts on day 14.

**Fig. 3:**
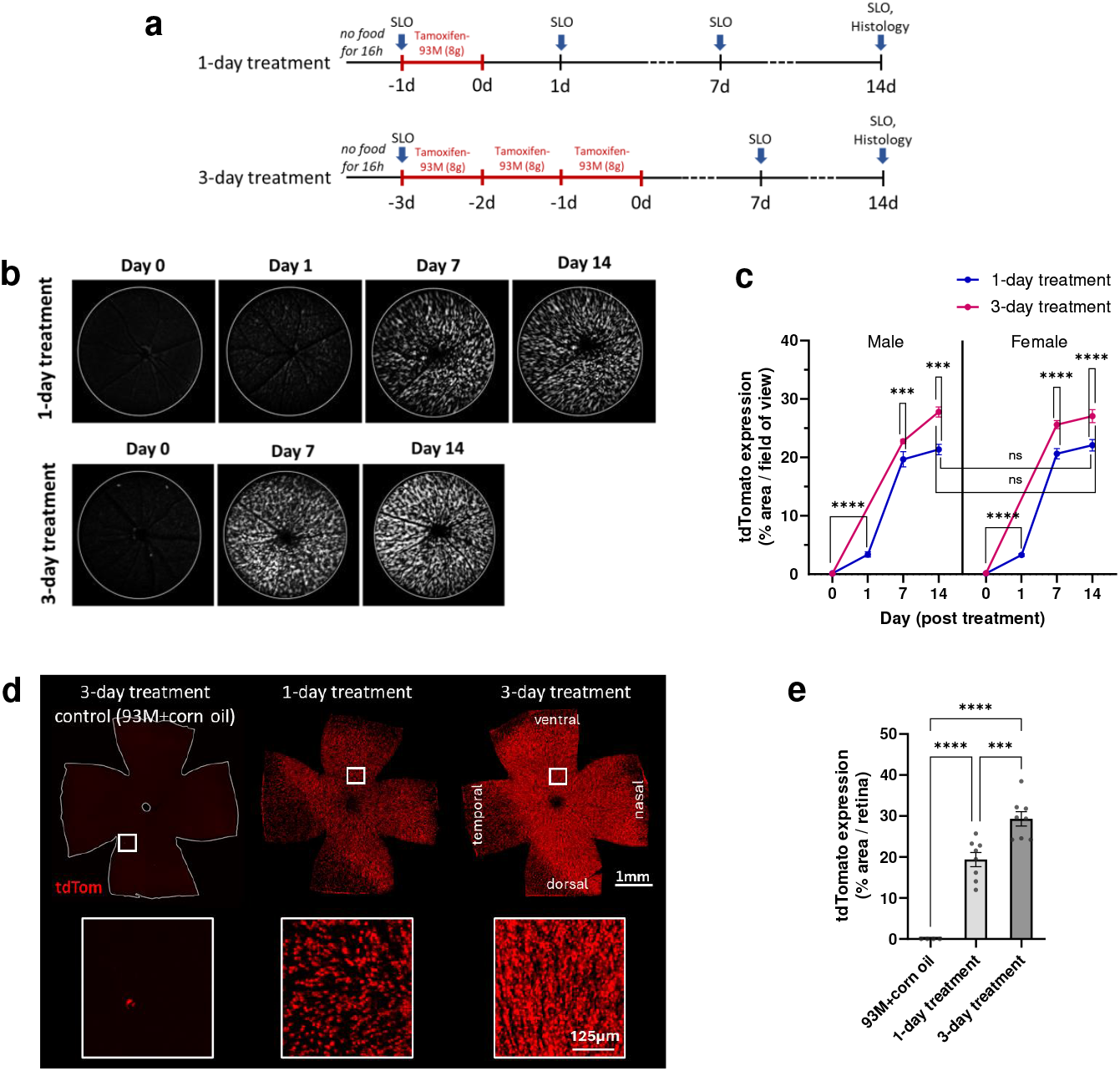
Feasibility and efficacy evaluation of Tamoxifen-infused 93M for Cre recombination using a mouse line with inducible tdTomato expression in retinal Müller glia. **a**, Experimental design for 1-day and 3-day Tamoxifen-93M treatment groups in Rlbp1-Cre^ERT2^;*Rosa*^*ai14*^ mice. Both groups received 8 g/d with an equal Tamoxifen concentration of 312.5 μg/g. The body weight and unconsumed Tamoxifen-93M were measured after each feeding cycle. SLO was performed for the 1-day treatment on day -1, 0, 1, 7, and 14, and on day -3, 0, 7, and 14 for the 3-day treatment. **b**, Representative longitudinal SLO image series per treatment group shows tdTomato expression induction *in vivo*. **c**, Quantification of the TdTomato expression as an area fraction of positive pixels in SLO images per sex and treatment group. **d**, Representative endpoint histology (day 14) of whole mounted retina per treatment group, including a control retina treated with corn oil-infused 93M. The reduced tdTomato expression in the dorso-temporal area is a commonly observed feature in the Rlbp1-Cre^ERT2^;*Rosa*^*ai14*^ line upon Tamoxifen treatment and therefore not specific to the diet gel-based treatment approach. **e**, Quantification of tdTomato expression as an area fraction of positive pixels in retinal whole mounts over both sexes per treatment group. N = 8 mice (4 males, 4 females) per Tamoxifen treatment group. Statistics: Two-way ANOVA with Tukey’s multiple comparisons test for (**c**) and one-way ANOVA with Tukey’s multiple comparisons test for (**e**). Results are shown as Mean ± SEM. Significance levels: \*\*\**P* < .001, \*\*\*\**P* < .0001.

SLO confirmed successful Tamoxifen delivery and Cre recombination, as shown by the induction of tdTomato expression over time (**Fig. 3b**). Interestingly, the 1-day treatment showed a significant increase (*P* < 0.0001) in the tdTomato expression for males and females after 24 hours, with a relative expression increase from 0.12% ± 0.04% to 3.39% ± 0.45% and 0.14% ± 0.03% to 3.29% ± 0.21%, respectively (**Fig. 3c**). The expression continued to increase until the end of the experiment on day 14 to 21.35% ± 0.88% for the males, and to 22.09% ± 0.98% for the females, without a significant difference between both sexes. The 3-day treatment on day 14 caused a small but significant increase in tdTomato expression compared to the 1-day treatment with a relative value of 27.78% ± 0.86% for the males (*P* = 0.007) and 27.05% ± 1.12%for the females (*P* < 0.0001) (**Fig. 3c**). No significant difference was observed between both sexes within the same treatment regimes.

The endpoint evaluation of the expression level in the whole mounted retina supports the treatment-time-dependent effect shown by the *in vivo* findings (**Fig. 3d**,**e**). The calculated overall tdTomato expression for males and females following the 1-day treatment is 19.38% ± 1.72% and 29.31% ± 1.76% for the 3-day treatment, with a significant difference between treatment times (*P* = 0.0010).

To control for potential effects caused by the corn oil (drug vehicle) in 93M, a separate SLO study was conducted using the 3-day treatment paradigm with corn oil-infused 93M (**Supplementary Fig. 3a**). The mice were imaged pre-treatment (baseline) and at days 7 and 14 post-treatment. The evaluation of the tdTomato expression *in vivo* showed no significant differences at both time points when compared to the baseline conditions, with a relative expression of 0.13% ± 0.04% at baseline, and 0.09% ± 0.03% and 0.10% ± 0.01% on day 7 and 14, respectively. In comparison, the endpoint evaluation of the whole mounted retina showed an average expression of 0.02% ± 0.01% (**Fig. 3e**).

Unlike for the PLX5622-93M feeding, Tamoxifen in 93M caused variations in the consumption rate between animals, with a tendency for females to leave more unconsumed Tamoxifen-93M after each feeding cycle (**Supplementary Fig. 3b**), but without significant body weight changes during the short treatment periods in comparison to the post-fasting conditions (**Supplementary Fig. 3c,d**). Since the difference in consumption could result in differences in Tamoxifen dosages between animals, a dose-response curve was calculated using a simple linear regression analysis of the total dose for each animal versus the tdTomato expression present in the retinal whole mounts on day 14 (**Fig. 4**). The total Tamoxifen dose per mouse (μg/g body weight) was calculated by the total amount (g) of consumed Tamoxifen-93M and the averaged body weights over the treatment time (1 or 3 days). Both treatment groups show a significant correlation between the dose and the tdTomato expression with an R^2^ of 0.5686 for the 1-day treatment and R^2^ of 0.6430 for the 3-day treatment.

**Fig. 4:**
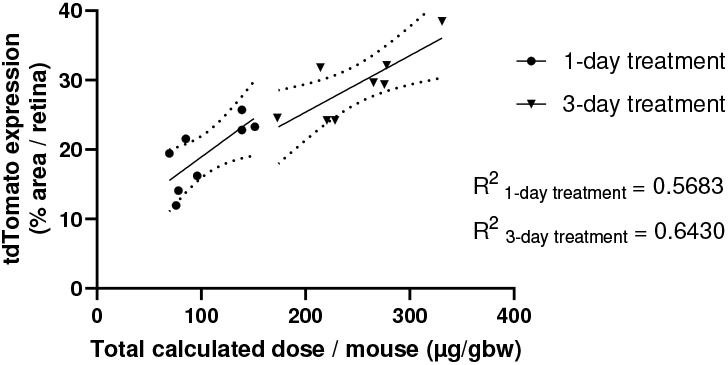
Dose-response curve of Tamoxifen-infused 93M. Linear regression analysis of the calculated total dose of Tamoxifen for each animal (μg/g body weight) versus the tdTomato expression level present in the retinal whole mounts of the 1-day and 3-day treatment groups on day 14. The total Tamoxifen dose per mouse (μg/g body weight) was calculated based on the total amount (g) of consumed Tamoxifen-93M and the averaged body weights over the treatment time (1 or 3 days). N = 16 mice. R^2^ for both treatment groups and 95% confidence intervals are shown.

## Discussion

Gel-based diets are often used to supplement chow for breeding purposes or in post-operative or compromised animals to prevent body weight loss ^22–28^. However, gel-based maintenance diets can also be used to fully replace chow ^29,30^. In this study, we identified that daily feedings of 8 g of 93M per mouse is a suitable amount that can be completely consumed and allows for steady body weight over time in adult mice of both sexes. Feeding a lower amount, namely 6 g/d, caused a steady drop in body weight which was more pronounced in males. This may be attributed to the higher average body weight of males with a mean difference between the sexes of 32.8% ± 2.1% on post-fast day 0 (*P* < 0.0001, unpaired two-tailed t-test). On the other hand, the 93M consumption data of age-matched mice indicates that females in the 10 g/d group tend, even though not significantly, to leave more unconsumed food after each feeding cycle than their male counterparts.

Furthermore, we show that the gel texture and the liquefication properties of 93M upon heating allow for a simple way to infuse water-insoluble drugs into 93M. Here, we tested the two small molecules, PLX5622 and Tamoxifen, with good results, and it is likely that this delivery system could work for other water-insoluble drugs or test compounds.

Unexpectedly, our PLX5622 experiment using the high dose of 2.0 mg/g revealed a significant dose-dependent effect on the body weights in male mice. This is a new and interesting observation that may imply sex-specific metabolic changes in males upon high-dose PLX5622 treatment, especially since PLX5622 has been shown to affect glucose metabolism and adipose tissue remodeling ^31,32^. This should be considered for studies in which a comparable dose is planned.

There are several advantages of using PLX5622-infused 93M for microglia depletion. This approach enables the use of different drug concentrations with consistent consumption rates over time in age-matched mice of both sexes. Another is the ability to easily and cost-effectively adjust dosage in small batches of food. These features make it practical to test PLX5622 from different vendors or between different production lots and adjust dosages accordingly, which is cost-prohibitive with chow-based preparations. Another advantage is the lower sucrose content of the AIN-93-based diet used here, which mainly uses corn starch as a carbohydrate source, compared to the commonly used high sucrose-based AIN-76A chow for PLX5622 and other drugs ^33,34^. In addition, the presented drug delivery system avoids the risks and animal welfare concerns of invasive methods of PLX5622 applications, such as i.p. injections of PLX5622-DMSO mixtures, which require daily injections ^35,36^. This method may be suitable for bigger rodents, such as Sprague Dawley rats, but we observed abdominal cyst formations during a trial of repeated PLX5622-DMSO i.p. injections into mice following a published protocol ^35^. Overall, the diet gel-based delivery of PLX5622 is highly efficient, mitigates adverse effects of other treatment routes, and allows for long-term exposure experiments or the testing of different batches of drugs with adjusted dosages.

We also demonstrate that Tamoxifen can be infused into 93M and that feeding it to adult and sex-matched mice induces Cre recombination. Here, a single Tamoxifen concentration (312.5 μg/g) and a daily feeding amount (8 g/day) were used while controlling for the treatment time (1-day vs. 3-day treatment). The longitudinal SLO data observing Cre-dependent tdTomato expression in retinal Müller glia shows that Tamoxifen works quickly, with tdTomato activation within 24 hours after one feeding cycle. Furthermore, the control over treatment time is sufficient to significantly adjust the level of Cre recombination, despite the differences in consumption between animals. An alternative to better control the level of Cre recombination may be adjusting the Tamoxifen concentration in 93M to the individual body weights of the mice. However, there might be a limitation on the amount of Tamoxifen that can be incorporated due to palatability. Our data shows that mice show some reluctance to Tamoxifen-infused 93M, which is more pronounced in females compared to male mice. Interestingly, studies using a commercially available Tamoxifen-incorporated chow reported similar observations with reduced feeding and weight loss within the first days of treatment ^37–39^. In our study, we only observed a limited drop in body weight over the short treatment periods of one and three days relative to the preceding post-fasting body weight (**Supplementary Fig. 3c,d**). Interestingly, mice treated with 93M or corn oil-infused 93M also showed a limited initial weight loss (**Fig. 1c,d** and **Supplementary Fig. 4a**), while this was not observed in the PLX5622-93M or DMSO-infused 93M treated mice (**Supplementary Fig.2c**,**d**, and **Supplementary Fig. 4b**) Thus, it remains unclear whether Tamoxifen, the switch to a gel-based diet, the mouse strain, or a combination of these factors contributes to the weight loss. Overall, a high level of recombination can be achieved with a repeated feeding regimen, whereas a single feeding can be advantageous for mosaic recombination. Importantly, Tamoxifen-infused 93M offers a refinement over gavage- or injection-based delivery.

In conclusion, our study shows that a gel-based diet such as 93M can be used as an alternative system for dosage-controlled drug delivery. It shows the advantage of staying within the balance of animal welfare concerns by minimizing injury, pain, or discomfort, and largely staying within the boundaries of gaining or losing body weight of ± 25 percent, which could be reduced by shortening fasting times. Furthermore, it allows for the preparation of small batches in the lab, and therefore provides the ability to test or titrate new batches of drug. This diet gel-based oral drug delivery system may also be expanded to other drugs that are water soluble or water insoluble.

### Limitations of approach

93M liquification requires heating at 60°C which could inactivate heat-labile molecules during incorporation. Another limitation may be the difficulty of increasing dosages in 93M if the animals avoid the drug-infused diet due to palatability issues.

### Limitations of study

The consumption behavior was not recorded. Differences between animals could change the outcomes depending on how quickly or slowly the animals eat the drug-infused 93M. Furthermore, the optimal daily consumption amount of 93M was defined for 8–12-week-old mice and may differ for other age groups. Another limitation is that we only investigated the effect of drug-infused 93 M in the retina using one Cre line and a microglia reporter line. The effectiveness of the drugs could differ in other tissues or in combination with different Cre or reporter lines.

## Methods

### Animals

For all experiments, mice were single-caged, and sex- and age-matched (between 8 and 12 weeks). Mice were maintained in a temperature and humidity-controlled animal facility in individually ventilated cages on a 12-hr light–dark schedule. All animals had access to food according to the experimental design and to water ad libitum. Prior to each experiment mice were deprived of food for 16 hours to increase appetite for a timely start of the experiment across all experimental animals and to reduce eventual reluctance of 93M consumption due to taste changes.

The study included three mouse strains: B6129SF1/J mice (RRID:IMSR_JAX:101043; The Jackson Laboratory, ME, USA), which were used for the evaluation of the optimal daily consumption rate of 93M. The *Cx3cr1*^*gfp/+*^ microglia reporter mice (B6.129P2(Cg)-*Cx3cr1* ^*tm1Litt*^/J; RRID:IMSR_JAX:005582, The Jackson Laboratory, ME, USA) were used for the PLX5622 study. The Rlbp1-Cre^ERT2^;*Rosa*^*ai14*^ mice (Gt(ROSA)26Sor^tm14(CAG-tdTomato)Hze^ Tg(Rlbp1-cre/ERT2)1Eml/Eml; RRID:MGI:7708085; Levine Lab, VUMC, TN, USA) were used for the Tamoxifen study ^19,20,40^. This study was approved by the Vanderbilt University Medical Center Institutional Animal Care and Use Committee and conformed to the Association for Research in Vision and Ophthalmology Statement for the Use of Animals in Ophthalmic and Vision Research.

### Experimental groups

Three experiments were designed. The first experiment evaluated the optimal daily consumption rate of 93M to ensure complete consumption while maintaining body weight by testing a low (6 g), an intermediate (8 g), and a high amount (10 g) of 93M provided daily in single-caged mice. The second tested the feasibility of PLX5622-infused 93M feeding as a dosage-controllable method to deplete microglia at a low (0.8 mg/g) and a high (2.0 mg/g) concentration. The third experiment tested whether Tamoxifen can be administered through 93M feeding at a single concentration (312.5μg/g) while controlling for the degrees of Cre recombination by adjusting the duration of the treatment (1 versus 3-day treatment).

### DietGel^®^

Three different DietGel^®^ products provided by ClearH_2_O^®^, Inc., ME, USA were tested for drug-vehicle incorporation, which included DietGel^®^ Boost, DietGel^®^ 93M, and DietGel^®^ GEM.

DietGel^®^ 93M (93M), a complete maintenance diet with enhanced flavor to promote consumption (https://clearh2o.com/products/dietgel%C2%AE-93m, accessed on November 26^th^, 2024), was selected for the study based on smoothness, liquefication properties upon heating, and emulsion properties of oil-based additives.

### PLX5622 and Tamoxifen incorporation into DietGel^®^ 93M

PLX5622 (C-1521, Lot#21; Chemgood, LLC., VA, USA) and Tamoxifen (T5648, Sigma-Aldrich, MO, USA) were separately incorporated into 93M per cup. Each cup contains 75g of 93M (measured in our lab).

PLX5622 was incorporated at two different concentrations to a final concentration of 0.8 mg/g and 2.0 mg/g. For the initial concentration, 60 mg of PLX5622 and 150 mg for the latter were each mixed into 1 ml of 100% DMSO (pharma grade, Heiltropfen Lab. LLP, UK) using a 1.5 ml microcentrifuge tube. The tubes were protected from light and placed on a rocker for constant rocking for 1 hour at room temperature, followed by a 20-minute ultrasonic bath to achieve complete dissolvement.

Tamoxifen was used at a concentration of 312.5 μg/g. For one cup of 93M, 23.44 mg of Tamoxifen was dissolved in 1 ml of corn oil (C8267, Sigma-Aldrich, MO, USA) on a rocker at room temperature overnight (light protected).

Each cup of 93M was heated in a water bath at 60°C for 15 minutes to liquify 93M. 1ml of the freshly prepared drug-vehicle solution was stirred in dropwise over 5 minutes using a P1000 pipet and a metal spatula, followed by 5 min of stirring to achieve a homogenous mixture. The cups were sealed with parafilm and placed on ice for 10 minutes before light-protected storage at 4°C. The dug-infused 93M can be stored for at least up to one week at 4°C.

To measure out the desired amount of DietGel^®^ for the experiment, a wooden single-use spatula was used to transfer it into a 5 cm glass petri dish, which was then placed into the mouse cage. The dishes were thoroughly cleaned out daily with paper towels before new food was added.

### Anesthesia and scanning laser ophthalmoscopy (SLO)

The custom-built multimodal MURIN system, including a 2-channel fluorescence SLO, was used for *in vivo* retinal fluorescence imaging ^41^. Mice were anesthetized using a mixture of isoflurane and oxygen set at a ratio of 2.5% and a flow rate of 1.5 l/min. Mydriasis was induced by applying 1-2 drops of 1% Tropicamide eye drops (VUMC pharmacy, Nashville, TN, USA).

The eyes were fitted with a custom zero-diopter contact lens to reduce optical aberrations at the air-cornea interface ^41^. During imaging, 0.3% hydroxypropyl methylcellulose lubricant eye gel (GenTeal^®^ Tears; Alcon) was periodically applied to preserve corneal hydration. Animals were placed onto a custom water-heated imaging bed maintained at 38°C with an integrated palate bar and nose cone to maintain anesthesia and reduce movement during the SLO imaging session. The green channel was used to image the GFP signal in the microglia-reporter mice B6.129P2(Cg)-Cx3cr1 ^*tm1Litt*^/J, and the red channel for the tdTomato expression in Müller glia of the Rlbp1-Cre^ERT2^;*Rosa*^*ai14*^ mice. Each SLO scan consisted of 200 acquired frames. Raw images were pre-processed as described elsewhere ^41^.

### Quantification of longitudinal microglia counts and relative tdTomato expression *in vivo* in SLO scans

Pre-processed SLO images were loaded into Fiji (ImageJ 1.54f, National Institutes of Health) and twenty to thirty consecutive frames with minimal motion were selected for averaged intensity orthogonal projection. The projection image was converted to an 8-bit image followed by background subtraction with the rolling bar radius set at 20 pixels. The images were further processed for noise reduction and therefore sharpening using the unsharp mask function with a radius set at 6 pixels and a mask weight of 0.6 for the green channel (GFP) and 0.3 for the red channel (tdTomato), respectively ^42,43^. A threshold was applied using the threshold function ‘Moments’. If needed, the threshold was manually adjusted to eliminate background noise. The ROI manager was used to outline the entire field of view, and the area fraction of positive pixels (% area) was automatically measured using the ‘Measure’ function after selecting ‘Limit to threshold’ and ‘Area fraction’ in the measurement settings. The area fraction is the relative tdTomato expression.

### Retinal whole mount preparation and fluorescence microscopy

All mice were euthanized at the end of each experiment for endpoint histology using CO_2_ asphyxiation and secondary cervical dislocation. Both eyes were dissected using ophthalmic micro-scissors and rinsed in 1x PBS. The eyes were fixed in 4% PFA overnight at 4°C. The left eye was used for retinal whole-mount preparation. In brief, the cornea and lens were removed using ophthalmic micro-scissors. The sclera along with the choroid and RPE were cut radially and peeled off to expose the retina and subsequently separated by cutting the optic nerve at its head. Next, four radial incisions were made into the retina before mounting on a glass slide with the ganglion cell layer facing up. The whole mounts were coversliped using Fluoromount-G™ Mounting Medium (Invitrogen) and stored protected from light at 4°C upon imaging.

The retinal whole mounts were scanned with a ZEISS Axio Zoom.V16 with Apotome 3 microscope at 40X digital magnification using a 1x objective. A 4×4 tile scan was performed to cover the entirety of the retina with a Z-stack interval set at 9 μm. The scanning depth was individually adjusted to include all signal-positive layers. Apotome images were created using the software’s internal image processing function, and tiles were fused with an overlap set at 20%.

### Endpoint quantification of microglia and relative tdTomato expression in retinal whole mounts

Apotome images were loaded into Fiji (ImageJ 1.54f, National Institutes of Health) for maximum intensity orthogonal projection. The projection image was converted to an 8-bit grayscale image. The background was subtracted with the rolling bar radius set at 50 pixels. A retinal ROI (rROI) outlining the retina without including the optical nerve head was created and saved by subtracting the optic nerve head area from the retinal area using the XOR function in the ROI manager. Next, separate image analysis protocols were used for images showing GFP-expressing microglia and tdTomato expressing Müller glia.

Microglia reporter images were filtered using the morphological filtering operation ‘Top-hat’ with a radius set at 3 pixels to homogenize the background and enhance the morphological boundaries (contrast enhancement) for accurate automated cell counting ^44–47^. A threshold was defined using the ‘Otsu’ settings and manually adjusted if needed. The ‘Analyze particles’ function was used for automated counting of individual cell bodies within the rROI. To do so, the size (micron^2) was set to 10 – infinity, and ‘Count Masks’ and ‘Add to Manager’ were selected. The output files included ROIs of each counted cell. The new ROIs were overlayed onto the original image to confirm proper cell counting. If inaccurate, the counting was repeated with an adjusted threshold.

For the images showing tdTomato expressing whole mounts, a threshold was defined using the function ‘Moments’ and, if necessary, manually adjusted to improve the background-to-signal ratio. To confirm proper thresholding, the function ‘Create Selection’ was used to create an ROI outlining all positive pixels, which was then pasted onto the original image. The threshold was adjusted if the selection was deemed inaccurate. The rROI was applied to the threshold image before the ‘Measure’ function was used to calculate the area fraction (% area) of tdTomato positive pixels within the rROI. The area fraction is the relative tdTomato expression in the retinal whole mounts.

### Statistics

The GraphPad Prism 10.4.1 software (GraphPad Software, Inc., La Jolla, CA, USA) was used for data visualization and statistical analysis. Results are shown as Mean ± SEM. A *P*-value below 0.05 was considered significant.

## Data availability

The data generated during the current study are available from the corresponding authors upon request.

## Acknowledgments

We thank the members of the Levine and Fuhrmann laboratories for their insights and support during the course of this project.

## References

1. Turner, P. V, Pekow, C., Vasbinder, M. A. & Brabb, T. Administration of substances to laboratory animals: equipment considerations, vehicle selection, and solute preparation. J Am Assoc Lab Anim Sci 50, 614–27 (2011).

2. Gad, S. C. et al. Tolerable Levels of Nonclinical Vehicles and Formulations Used in Studies by Multiple Routes in Multiple Species With Notes on Methods to Improve Utility. Int J Toxicol 35, 95–178 (2016).

3. Anselmo, A. C., Gokarn, Y. & Mitragotri, S. Non-invasive delivery strategies for biologics. Nat Rev Drug Discov 18, 19–40 (2019).

4. Jain, K. K. An Overview of Drug Delivery Systems. in Methods in molecular biology (Clifton, N.J.) vol. 2059 1–54 (2020).

5. Hovard, A., Teilmann, A., Hau, J. & Abelson, K. The applicability of a gel delivery system for self-administration of buprenorphine to laboratory mice. Lab Anim 49, 40–45 (2015).

6. Overk, C. R., Borgia, J. A. & Mufson, E. J. A novel approach for long-term oral drug administration in animal research. J Neurosci Methods 195, 194–199 (2011).

7. Zhong, L. et al. Small molecules in targeted cancer therapy: advances, challenges, and future perspectives. Signal Transduct Target Ther 6, 201 (2021).

8. Beck, H., Härter, M., Haß, B., Schmeck, C. & Baerfacker, L. Small molecules and their impact in drug discovery: A perspective on the occasion of the 125th anniversary of the Bayer Chemical Research Laboratory. Drug Discov Today 27, 1560–1574 (2022).

9. Spangenberg, E. et al. Sustained microglial depletion with CSF1R inhibitor impairs parenchymal plaque development in an Alzheimer’s disease model. Nat Commun 10, 3758 (2019).

10. Ebneter, A., Kokona, D., Jovanovic, J. & Zinkernagel, M. S. Dramatic Effect of Oral CSF-1R Kinase Inhibitor on Retinal Microglia Revealed by In Vivo Scanning Laser Ophthalmoscopy. Transl Vis Sci Technol 6, 10 (2017).

11. Jovanovic, J., Liu, X., Kokona, D., Zinkernagel, M. S. & Ebneter, A. Inhibition of inflammatory cells delays retinal degeneration in experimental retinal vein occlusion in mice. Glia 68, 574–588 (2020).

12. Wickel, J. et al. Repopulated microglia after pharmacological depletion decrease dendritic spine density in adult mouse brain. Glia 72, 1484–1500 (2024).

13. Park, E. J. et al. System for tamoxifen-inducible expression of cre-recombinase from the Foxa2 locus in mice. Developmental Dynamics 237, 447–453 (2008).

14. Jardí, F. et al. A shortened tamoxifen induction scheme to induce CreER recombinase without side effects on the male mouse skeleton. Mol Cell Endocrinol 452, 57–63 (2017).

15. Elmore, M. R. P. et al. Colony-Stimulating Factor 1 Receptor Signaling Is Necessary for Microglia Viability, Unmasking a Microglia Progenitor Cell in the Adult Brain. Neuron 82, 380–397 (2014).

16. Ichise, H. et al. Establishment of a tamoxifen-inducible Cre-driver mouse strain for widespread and temporal genetic modification in adult mice. Exp Anim 65, 231–244 (2016).

17. Navabpour, S., Kwapis, J. L. & Jarome, T. J. A neuroscientist’s guide to transgenic mice and other genetic tools. Neurosci Biobehav Rev 108, 732–748 (2020).

18. Chen, M.-Y., Zhao, F.-L., Chu, W.-L., Bai, M.-R. & Zhang, D.-M. A review of tamoxifen administration regimen optimization for Cre/loxp system in mouse bone study. Biomedicine & Pharmacotherapy 165, 115045 (2023).

19. Stone, M. L., Lee, H. H. & Levine, E. M. Agarose hydrogel-mediated electroporation method for retinal tissue cultured at the air-liquid interface. iScience 27, 111299 (2024).

20. Pollak, J. et al. ASCL1 reprograms mouse Müller glia into neurogenic retinal progenitors. Development 140, 2619–2631 (2013).

21. Webster, M. K. et al. Stimulation of Retinal Pigment Epithelium With an α7 nAChR Agonist Leads to Müller Glia Dependent Neurogenesis in the Adult Mammalian Retina. Investigative Opthalmology & Visual Science 60, 570 (2019).

22. Felgenhauer, J. L. et al. Evaluation of Nutritional Gel Supplementation in C57BL/6J Mice Infected with Mouse-Adapted Influenza A/PR/8/34 Virus. Comp Med 70, 471–486 (2020).

23. Gates, K. V, Alamaw, E., Jampachaisri, K., Huss, M. K. & Pacharinsak, C. Efficacy of Supplemental Diet Gels for Preventing Postoperative Weight Loss in Mice (Mus musculus). Journal of the American Association for Laboratory Animal Science 62, 87–91 (2023).

24. Wong, R. K. et al. Effects of Supplemental Diet during Breeding on Fertility, Litter Size, Survival Rate, and Weaning Weight in Mice (Mus musculus). Journal of the American Association for Laboratory Animal Science 63, 480–487 (2024).

25. Froberg-Fejko, K. M. & Lecker, J. Overview and indications for use of Bio-Serv’s Nutra-Gel diet for laboratory rodents. Lab Anim (NY) 40, 326–327 (2011).

26. Wong, R. K. et al. Effects of Supplemental Diet during Breeding on Fertility, Litter Size, Survival Rate, and Weaning Weight in Mice (Mus musculus). Journal of the American Association for Laboratory Animal Science 63, 480–487 (2024).

27. Young, M. S. et al. Subcutaneous Alfaxalone-Xylazine-Buprenorphine for Surgical Anesthesia and Echocardiographic Evaluation of Mice (Mus musculus). Journal of the American Association for Laboratory Animal Science 63, 49–56 (2024).

28. Boatman, S. et al. Diet-induced shifts in the gut microbiota influence anastomotic healing in a murine model of colonic surgery. Gut Microbes 15, (2023).

29. Imamura, Y. et al. Ultrasound stimulation of the vagal nerve improves acute septic encephalopathy in mice. Front Neurosci 17, (2023).

30. Iban-Arias, R. et al. Ad-derived bone marrow transplant induces proinflammatory immune peripheral mechanisms accompanied by decreased neuroplasticity and reduced gut microbiome diversity affecting AD-like phenotype in the absence of Aβ neuropathology. Brain Behav Immun 118, 252–272 (2024).

31. Bosch, A. J. T. et al. CSF1R inhibition with PLX5622 affects multiple immune cell compartments and induces tissue-specific metabolic effects in lean mice. Diabetologia 66, 2292–2306 (2023).

32. Ali, S. et al. CSF1R inhibitor PLX5622 and environmental enrichment additively improve metabolic outcomes in middle-aged female mice. Aging 12, 2101–2122 (2020).

33. Reeves, P. G., Nielsen, F. H. & Fahey, G. C. AIN-93 Purified Diets for Laboratory Rodents: Final Report of the American Institute of Nutrition Ad Hoc Writing Committee on the Reformulation of the AIN-76A Rodent Diet. J Nutr 123, 1939–1951 (1993).

34. Reeves, P. G. Components of the AIN-93 Diets as Improvements in the AIN-76A Diet. J Nutr 127, 838S–841S (1997).

35. Riquier, A. J. & Sollars, S. I. Astrocytic response to neural injury is larger during development than in adulthood and is not predicated upon the presence of microglia. Brain Behav Immun Health 1, 100010 (2020).

36. Riquier, A. J. & Sollars, S. I. Terminal field volume of the glossopharyngeal nerve in adult rats reverts to prepruning size following microglia depletion with PLX5622. Dev Neurobiol 82, 613–624 (2022).

37. Andersson, K. B., Winer, L. H., Mørk, H. K., Molkentin, J. D. & Jaisser, F. Tamoxifen administration routes and dosage for inducible Cre-mediated gene disruption in mouse hearts. Transgenic Res 19, 715–725 (2010).

38. Chiang, P.-M. et al. Deletion of TDP-43 down-regulates Tbc1d1, a gene linked to obesity, and alters body fat metabolism. Proceedings of the National Academy of Sciences 107, 16320–16324 (2010).

39. Miró-Murillo, M. et al. Acute Vhl gene inactivation induces cardiac HIF-dependent erythropoietin gene expression. PLoS One 6, e22589 (2011).

40. Vazquez-Chona, F. R., Clark, A. M. & Levine, E. M. Rlbp1 Promoter Drives Robust Mü ller Glial GFP Expression in Transgenic Mice. Investigative Opthalmology & Visual Science 50, 3996 (2009).

41. Rico-Jimenez, J. J. et al. MURIN: Multimodal Retinal Imaging and Navigated-laser-delivery for dynamic and longitudinal tracking of photodamage in murine models. Frontiers in Ophthalmology 3, (2023).

42. Penazzi, L., Sündermann, F., Bakota, L. & Brandt, R. Machine Learning to Evaluate Neuron Density in Brain Sections. in 263–291 (2014). doi:10.1007/978-1-4939-0381-8_13.

43. Ogura, A., Hayakawa, K., Miyati, T. & Maeda, F. Improvement on detectability of early ischemic changes for acute stroke using nonenhanced computed tomography: Effect of matrix size. Eur J Radiol 76, 162–166 (2010).

44. Li, J. et al. Segmentation of retinal microaneurysms in fluorescein fundus angiography images by a novel three-step model. Front Med (Lausanne) 11, (2024).

45. Bai, X., Zhou, F. & Xue, B. Image enhancement using multi scale image features extracted by top-hat transform. Opt Laser Technol 44, 328–336 (2012).

46. Bai, X., Zhou, F. & Xue, B. Toggle and top-hat based morphological contrast operators. Computers & Electrical Engineering 38, 1196–1204 (2012).

47. Ghislain, F., Beaudelaire, S. T. & Daniel, T. An accurate unsupervised extraction of retinal vasculature using curvelet transform and classical morphological operators. Comput Biol Med 178, 108801 (2024).

