## Supplementary Figures for "Diet gel-based oral drug delivery system for controlled dosing of small molecules for microglia depletion and inducible Cre recombination in mice"

**Supplementary Figure 1**

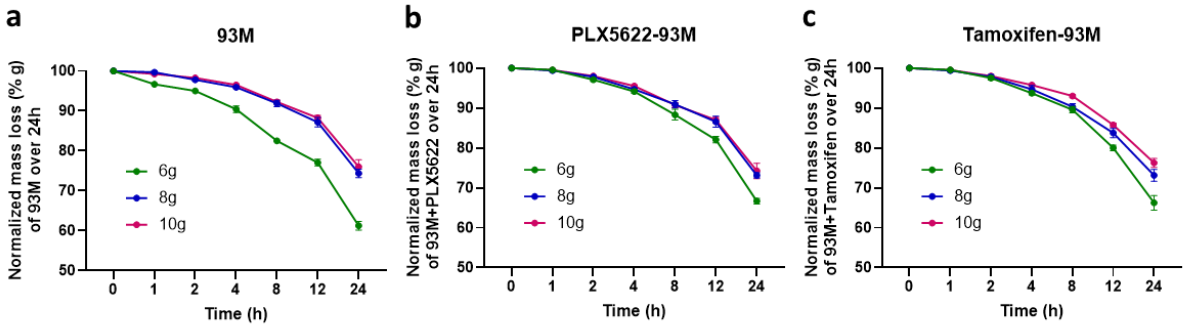


**Supplementary Fig. 1: Temporal mass reduction of 93M with and without drug infusion due to evaporation.** The different amounts of 93M (6 g, 8 g, 10 g) with and without drug infusions held in 5 cm glass petri dishes were placed individually into ventilated mouse cages. The weight was measured after 1, 2, 4, 8, 12 and 24 hours to establish mass reduction caused by evaporation. N = 4 per amount.

**Supplementary Figure 2**

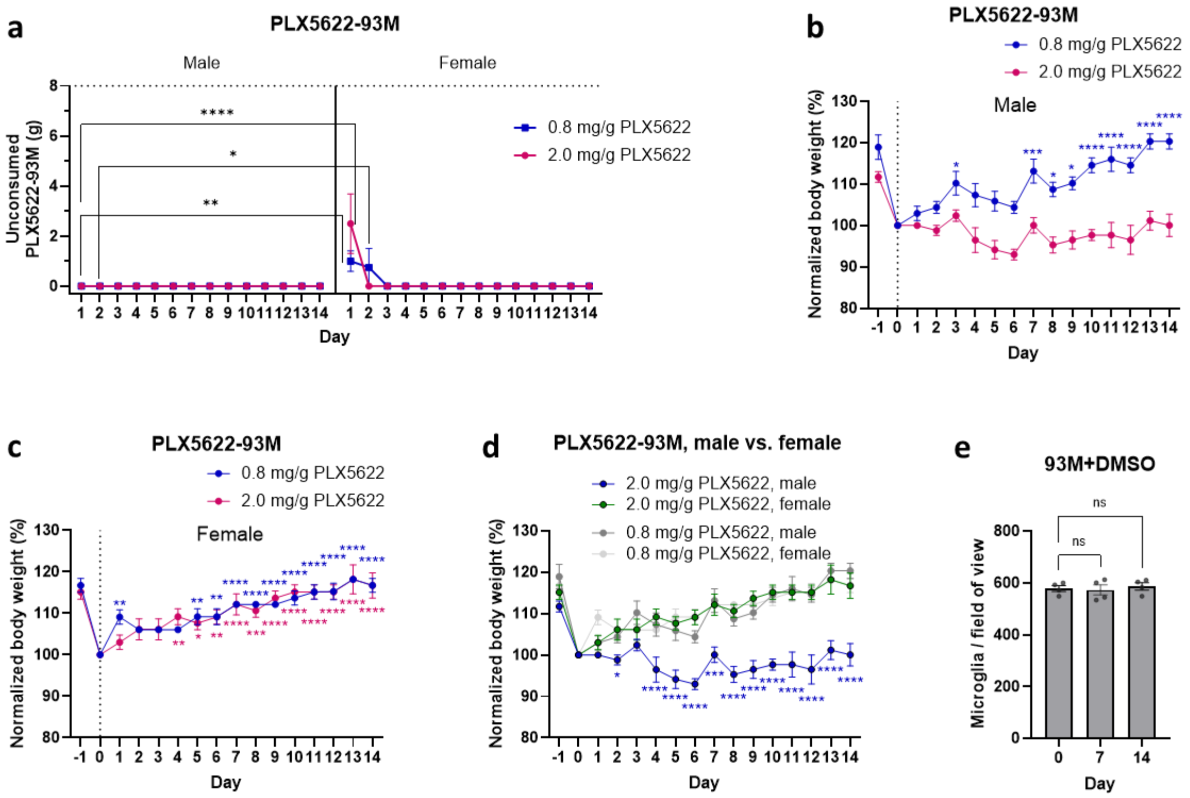


**Supplementary Fig. 2: Supplementary data of the PLX5622-infused 93M experiment. a**, The average mass of unconsumed PLX5622-infused 93M after each feeding cycle (24h) of the 0.8 mg/g and the 2.0 mg/g dosage group separated by sex. **b**, **c**, Temporal body weight measurements normalized and compared to the post-fasting body weight (day 0, dotted line) in males and females for both PLX5622-93M dosage groups. N = 8 mice (4 males, 4 females) per dosage group. **d**, Aggregation of the data from (**b** and **c**). The shown significance levels are for the males (2.0 mg/g) in comparison to the female group (2.0 mg/g) at each timepoint. These males also significantly differ from the two other groups from day 4 onwards. **e**, Temporal quantification of GFP+ microglia in heterozygous B6.129P2(Cg)-*Cx3cr1* $^{tm1Litt}$/J mice fed with 8 g/d of DMSO-infused 93M (vehicle-control) for 14 days using retinal *in vivo* imaging (SLO). N = 4 eyes. Statistics: Two-way ANOVA with Tukey's multiple comparisons test for (**a**), two-way ANOVAs with Dunnett's multiple comparisons tests for (**b**,**c**), two-way ANOVA with Tukey's multiple comparisons test for (**d**), and one-way ANOVA with Dunnett's multiple comparisons test for (**e**). Results are shown as Mean ± SEM. Significance levels: $*P < .05$, $**P < .01$, $***P < .001$, $****P < .0001$.

**Supplementary Figure 3**

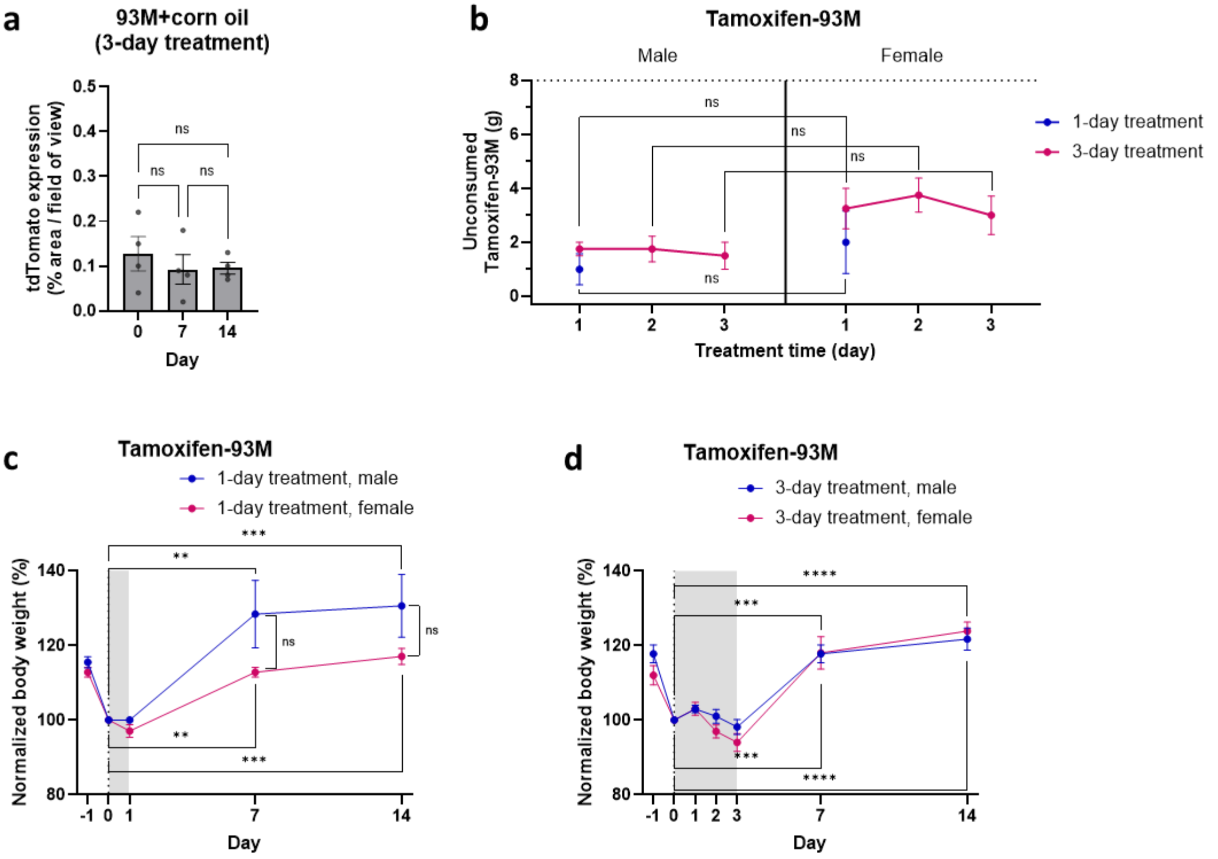


**Supplementary Fig. 3: Supplementary data of the Tamoxifen-infused 93M experiment. a**, Temporal quantification of the tdTomato expression in heterozygous Rlbp1-Cre$^{ERT2}$;*Rosa*$^{ai14}$ mice fed with 8 g/d of corn oil-infused 93M (vehicle-control) for 14 days using retinal *in vivo* imaging (SLO).N = 4 eyes. **b**, Averaged mass of unconsumed Tamoxifen-infused 93M after each feeding cycle (24h) of the 1-day and 3-day treatment group separated by sex. **c**,**d**, Temporal body weight measurements normalized and compared to the post-fasting body weight (day 0, dotted line) following the 1-day and the 3-day treatment with Tamoxifen-93M separated by sex. The gray shaded area indicates the treatment window of Tamoxifen-93M exposure. N = 8 mice (4 males, 4 females) per Tamoxifen treatment group. Statistics: One-way ANOVA with Tukey's multiple comparisons test for (**a**), two-way ANOVA with Tukey's multiple comparisons test for (**b**), and two-way ANOVA with Sidak's multiple comparisons tests for (**c**,**d**). Results are shown as Mean ± SEM. Significance level: $^{**}P < .01$, $^{***}P < .001$, $^{****}P < .0001$.

**Supplementary Figure 4**

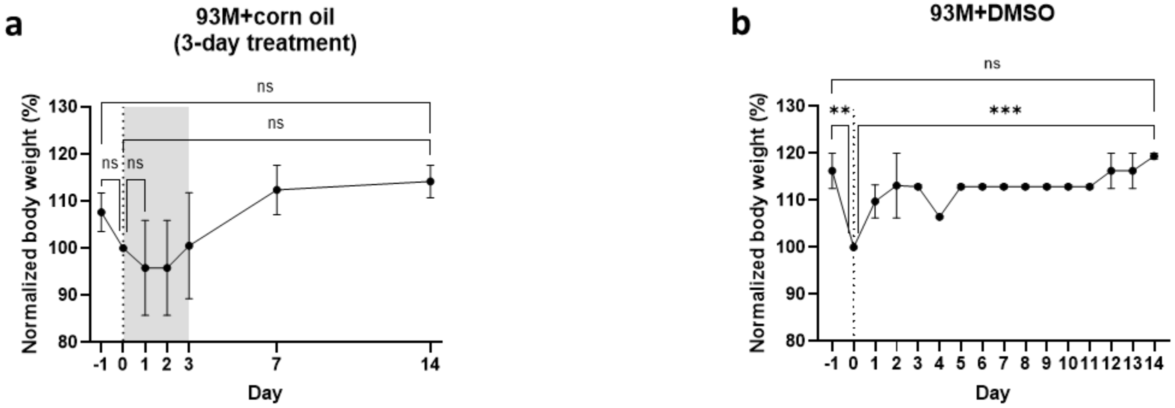


**Supplementary Fig. 4: Supplementary data of the body weight measurements in the vehicle-infused 93M feeding paradigms (vehicle control). a**,**b**, Temporal body weight measurements normalized and compared to post-fasting body weight (day 0, dotted line) in males and females for (**a**) corn oil-infused 93M and (**b**) DMSO-infused 93M. The Gray shaded area in (**a**) indicates the 3-day treatment window of Tamoxifen-93M exposure. N = 2 mice per vehicle-infused 93M evaluation. Statistics: Two-way ANOVA with Tukey's multiple comparisons test for (**a**,**b**). Results are shown as Mean ± SEM. Significance levels: **$P < .01$, ***$P < .001$.
